# Rapid preimplantation genetic screening (PGS) using a handheld, nanopore-based, DNA sequencer

**DOI:** 10.1101/274563

**Authors:** Shan Wei, Zachary R. Weiss, Pallavi Gaur, Eric Forman, Zev Williams

## Abstract

**Objective:** To determine if a handheld, nanopore-based DNA sequencer can be used for rapid preimplantation genetic screening (PGS).

**Design:** Retrospective study.

**Setting:** Academic medical center.

**Patient(s):** Amplified genomic DNA from euploid and aneuploid trophectoderm biopsy samples (n=9) that was also tested using traditional next generation sequencing (NGS).

**Intervention(s):** Short-read DNA library preparation and nanopore-based sequencing using a hand-held MinION sequencer.

**Main outcome measure(s):** Comparison of cytogenetic testing result from NGS and nanopore-based sequencing and the time required for library preparation and sequencing.

**Result(s):** Multiplexed short-read DNA library preparation was completed in 45 minutes. Sequencing times varied from 1 to 2 hours. These times compare favorably with NGS library preparation (>3.5 hours) and sequencing (>12 hours) times. Whole-chromosome aneuploidy screening results obtained from nanopore-based sequencing were identical to those obtained using NGS.

**Conclusion(s):** Methods for PGS of embryos have evolved from FISH to microarrays and most recently to NGS. Here we report the first application of nanopore-based sequencing for PGS on trophecoderm biopsy samples using a rapid multiplex short-read nanopore sequencing library preparation. Aneuploidy screening could be performed on 5 samples in one nanopore flowcell with 1 to 2 hour sequencing times. Overall, nanopore sequencing is a promising tool to perform rapid PGS assay onsite with a rapid turnover time, enabling same day testing and embryo transfer thus obviating the need for complex, large and expensive DNA sequencers or frozen embryos.

## Introduction

Human embryos have high rates of chromosomal aneuploidy, a significant cause of implantation failure and spontaneous pregnancy loss (1, 2). Preimplantation genetic screening (PGS) is an effective method to assess the numerical chromosomal constitution of embryos prior to transfer in patients undergoing IVF. PGS facilitates the selection of euploid embryos for transfer, thereby improving implantation rates and reducing the rates of miscarriage (3-5). By improving implantation rates in IVF, PGS empowers the practice of elective single embryo transfer (6, 7) thereby avoiding the increased obstetrical complications resulting from multiple gestation which occurs frequently after multiple embryo transfer.

Initially, PGS involved fluorescent in-situ hybridization (FISH) to assess the ploidy status of a subset of chromosomes. Likely due to limitations of the technology and the impact of blastomere biopsy at the cleavage stage, FISH-based PGS was shown to not be beneficial (8). Subsequent technologies included array-based comparative genomic hybridization (aCGH), single nucleotide polymorphism (SNP) microarray and quantitative real time polymerase chain reaction (qPCR) that enabled analysis of all chromosomes. These methods were limited by their high cost per sample and low throughput (9). Over the past several years, second-generation sequencing technologies – also known as next-generation sequencing (NGS) – consisting of massively-paralleled, high-throughput methods utilizing sequencing-by-synthesis-based chemistries have become increasingly widely used due to their higher dynamic range, lower cost, and ability to detect mosaicism (10-15). However, NGS-based aneuploidy detection still requires the use of complex and costly DNA sequencers and requires >12 hours for library preparation and sequencing, thereby necessitating embryo cryopreservation and often requiring the use of specialized reference laboratories.

Recently, third-generation sequencing technologies based on single-molecule sequencing have been developed. A prominent example is nanopore-based sequencing in which protein nanopores, approximately 10 nm in diameter, are distributed across a flow-cell membrane. An electrical current drives single-stranded DNA through the nanopore and the voltage changes that occur as each nucleotide passes through the nanopore are recorded. The identity of the nucleic acid is determined based on the variable resistivity of each base. Compared to traditional NGS technologies, nanopore-based sequencing has the distinct advantage of sequencing 15,000 times faster, delivering sequencing results in real-time and a much lower equipment cost.

The first commercial sequencer using nanopore technology is the MinION, which was released by Oxford Nanopore Technologies in 2014. The MinION is an 87gram hand-held device that is the size of a deck of cards and is powered by a USB cable attached to a computer. We developed a method for rapid library preparation and sequencing of short DNA fragments (cite: Genomics paper) and demonstrated that this method can provide rapid and aCGH-concordant aneuploidy detection from chorionic villus cells retrieved from products of miscarriage (16). We then developed a method to barcode and multiplex multiple samples within a single sequencing run (17). NGS technologies typically require days in order for results to become available, precluding a fresh embryo transfer when a biopsy is obtained at the blastocyst stage of development. The speed of the MinION based approach would provide PGS results within a few hours and therefore allow for potential fresh embryo transfer, avoiding the need for freeze-all cycles when PGS is employed.

In the present study, we apply our MinION rapid multiplexed short-read sequencing technique to aneuploidy detection in PGS and demonstrate that our optimized MinION library preparation and data analysis protocol is concordant to VeriSeq PGS aneuploidy results from DNA obtained after trophecoderm biopsy.

## Material and methods

### Samples

Trophectoderm 3-5 cell biopsy samples from fresh day 5 embryos were sent to a reference lab for routine clinical PGS testing. Samples were subjected to SurePlex whole genome amplification (WGA) and testing using the VeriSeq PGS assay following manufacturer’s instructions as part of routine clinical care. De-identified excess DNA was then sent to our research laboratory for of MinION library preparation and sequencing. 9 samples (n=9) were included in this retrospective study. The study was approved by the Institutional Review Board at Albert Einstein College of Medicine and Columbia University Medical Center.

### Library preparation

For each sample, ∼260ng WGA amplified products were also subjected to MinION PGS assay using rapid multiplex library preparation according to our recently developed protocol (Figure 1) (17). The full protocol is included in supplementary methods. Nine blinded samples consisting of diploid and aneuploid specimens, and one reference normal male sample, were tested using two MinION sequencing runs on the MinION MIN106 flowcell (Oxford Nanopore, MIN106).Each sequencing run consisted of one barcoded known reference sample and four blinded test samples. Initially a 2-hour sequencing run was performed using MINKNOW software version 1.7, and data were used for data analysis for aneuploidy detection. Additional 1-hour and 4-hour sequencing runs were performed after the initial sequencing run for mosaic aneuploidy and large CNV detection.

**Figure 1:**
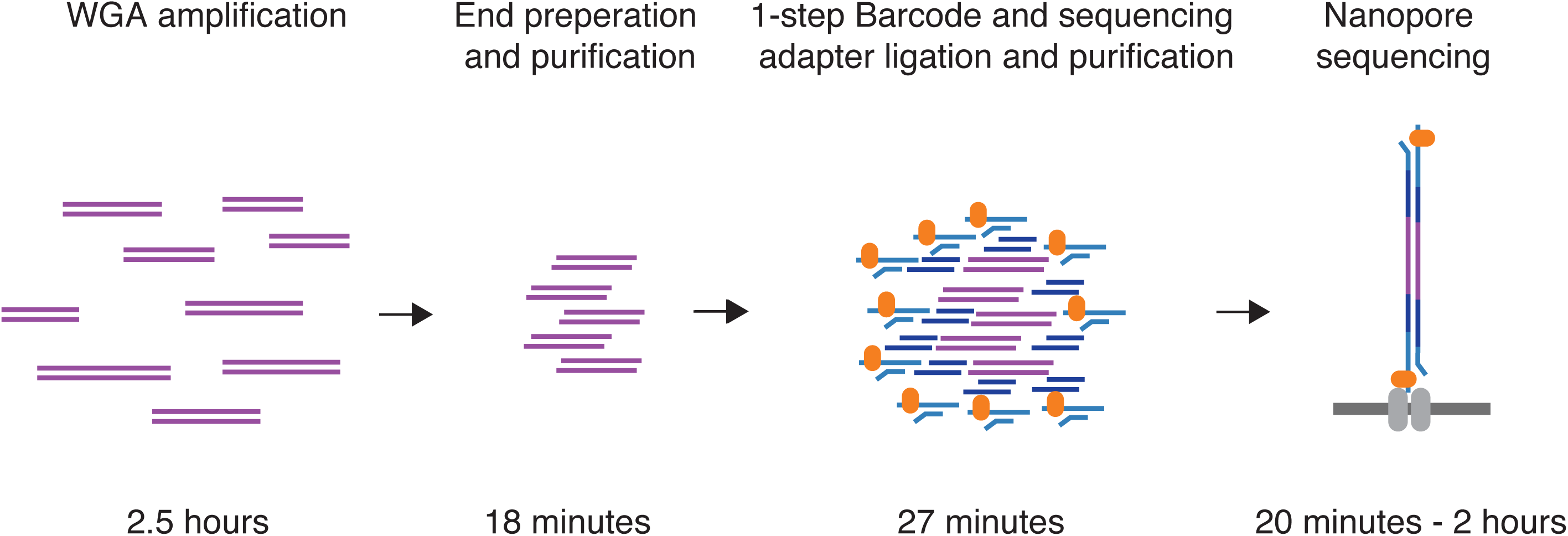
Illustration of library preparation for rapid nanopore sequencing. The workflow includes a standard ∼2.5-hour SurePlex whole genome amplification, ∼18 minute rapid end-preparation and purification, a 1-step barcode and sequencing adapter ligation and purification, and a 20-minute to 2-hour nanopore sequencing run, depending on the number of samples multiplexed into a single run.

### Data analysis

Sequences were translated locally to fastq from fast5 format by MINKNOW local basecaller and Albacore. Fastq format sequences were converted to fasta format and subjected to cutadapt v1.14 (18) for demultiplexing and short read removal using the parameter -O 20 -e 0.10 -m 50. The de-multiplexed sequences from each sample were subjected to parallel blat (pBlat) alignment to human reference genome GRCh37 and uniquely mapped reads were screened by pslReps from the UCSC suite (19) using parameter -minCover=0.40, -minAli=0.80, -nearTop=0.001and – singleHit. Numbers of sequences assigned to each chromosome were subjected to a modified Z-score method for aneuploidy detection as described in our former study (16, 17). Chromosomes with ≥ 25% copy number changes comparing to a normal male reference were considered not normal chromosomes, and they were eliminated in calculation of standard deviation for normal chromosomes (SD_normal) to increase the detection sensitivity for WGA PGS samples. Statistical analysis were performed in R v3.4.0 (20). Samples had been de-identified and the researcher was blinded to the VeriSeq PGS results at aneuploidy screening. A known normal male sample was included in each run as a reference.

For detection of CNV (≥ 20Mbs), 60,000 reads were used. Large CNVs were detected by segregating human genome reference into 10Mbs bins, and aneuploidy detection was performed for each bin as described above. Bins with fewer then 100 UA reads in the normal male reference samples were eliminated from large CNV detection assay. For bins with ≥ 200 UA reads in the reference sample, bins with Z-score > 3.29 were considered as gain on CNV, and Bins with Z-score < -3.29 was considered as loss in CNV (p=0.001); for bins with < 200 UA reads in the normal male reference sample, bins with Z-score > 6 were considered as gain on CNV, and bins with Z-score < 6 were considered as loss in CNV (p< 0.0001). Two bins with | Z-score | > 3.29 were concatenated. Large CNVs ≥ 20Mbs were reported.

60K UA reads were used for an investigation of possibility of mosaic aneuploidy detection. The mosaic aneuploidy in a 5-cell biopsy can range from 20-80%, and a mosaic aneuploidy in a 3-cell biopsy can range from 33-67%. Chromosomes with ≥ 15% copy number changes (∼30% mosaicism) comparing to a normal male reference were arbitrarily considered as mosaic aneuploidy in high confidence and not included in calculation of standard deviation of normal chromosomes (SD_normal) to increase detection sensitivity for WGA PGS samples.

## Results

### Sequencing performance

Two nanopore sequencing runs, each comprised of five barcoded samples, were performed. Samples included 3 normal samples (two normal males and one normal female); 2 samples with aneuploidy of a single chromosome (one female Monosomy 22, one female Trisomy 19), and 4 samples with aneuploidy on more then one chromosome (a female with Trisomy 22 and Monosomy X, a female with Trisomy 13 and Monosomy 14, a male with XXY and Trisomy 15, a female with Mosaic Trisomy 6, Trisomy 15 and Monosomy 18) (Table 1). Sample 6 consists of aneuploidy on Chr13 and Chr15 and a and large CNV gain on chr 15 (78,914,003-101,997,386) (Table 1). In Run 1, 74,001, 144,129, 288,489 and 548,166 reads were generated in 15, 30, 60 and 120 minutes, respectively. In Run 2, 89,027, 171,520, 347,397 and 664,769 reads were generated in 15, 30, 60 and 120 minutes, respectively (Figure 2A, Table 1). The rate of sequencing runs obtained 274K-332K reads per hour through the testing. Of reads generated, >70% were able to be assigned to a unique barcode. Of the assigned reads, 75-95% reads from each sample were uniquely assigned (UA) to the genome reference library (Hg19) (Table 1). Only those reads that were assigned to a barcode and uniquely matched to the reference genome library were used for subsequent downstream data analysis.

**Table 1.**
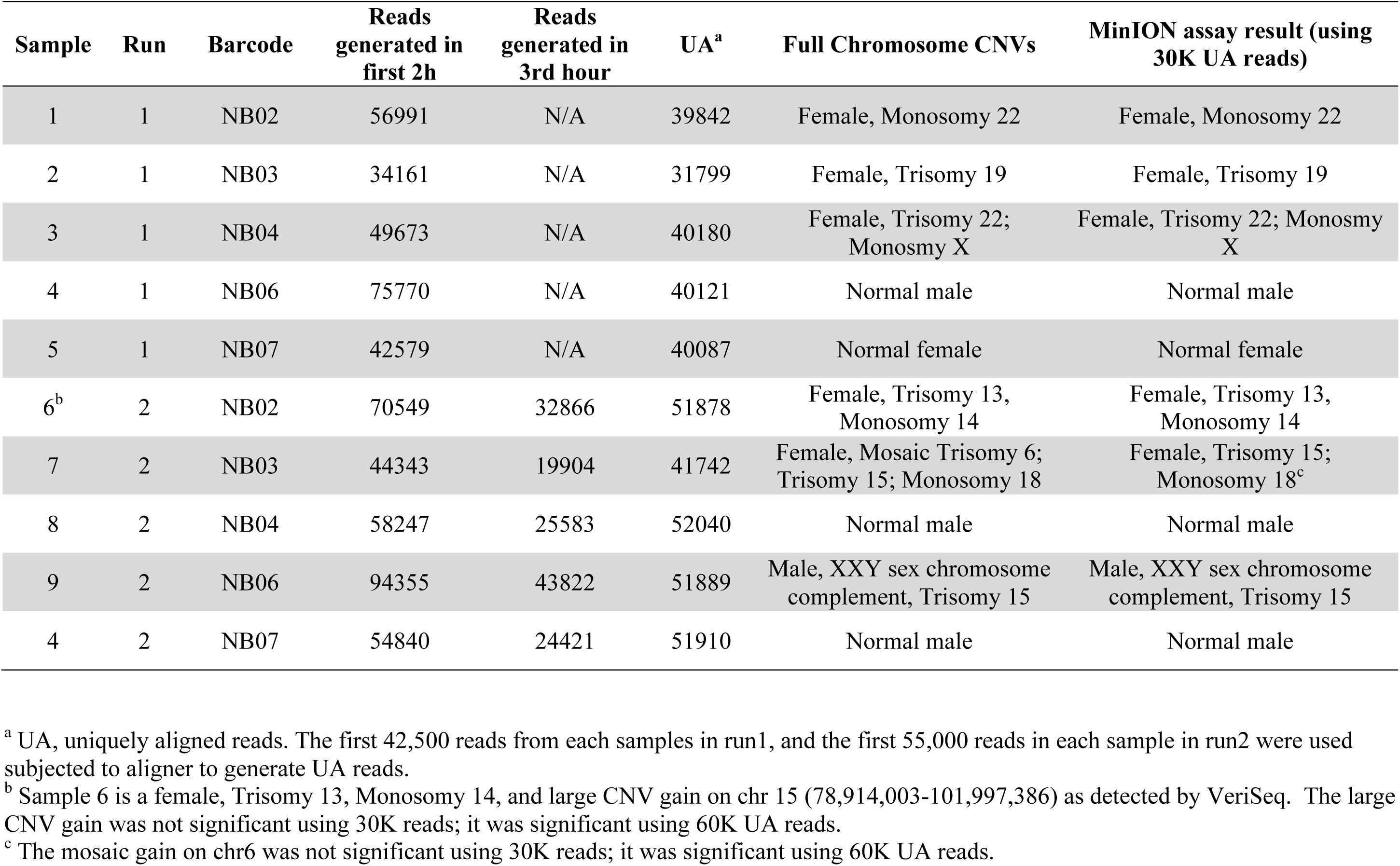
Run summary

| Sample | Run | Barcode | Reads generated in first 2h | Reads generated in 3rd hour | UA<sup>a</sup> | Full Chromosome CNVs | MinION assay result (using 30K UA reads) |
| --- | --- | --- | --- | --- | --- | --- | --- |
| 1 | 1 | NB02 | 56991 | N/A | 39842 | Female, Monosomy 22 | Female, Monosomy 22 |
| 2 | 1 | NB03 | 34161 | N/A | 31799 | Female, Trisomy 19 | Female, Trisomy 19 |
| 3 | 1 | NB04 | 49673 | N/A | 40180 | Female, Trisomy 22; Monosmy X | Female, Trisomy 22; Monosmy X |
| 4 | 1 | NB06 | 75770 | N/A | 40121 | Normal male | Normal male |
| 5 | 1 | NB07 | 42579 | N/A | 40087 | Normal female | Normal female |
| 6<sup>b</sup> | 2 | NB02 | 70549 | 32866 | 51878 | Female, Trisomy 13, Monosomy 14 | Female, Trisomy 13, Monosomy 14 |
| 7 | 2 | NB03 | 44343 | 19904 | 41742 | Female, Mosaic Trisomy 6; Trisomy 15; Monosomy 18 | Female, Trisomy 15; Monosomy 18<sup>c</sup> |
| 8 | 2 | NB04 | 58247 | 25583 | 52040 | Normal male | Normal male |
| 9 | 2 | NB06 | 94355 | 43822 | 51889 | Male, XXY sex chromosome complement, Trisomy 15 | Male, XXY sex chromosome complement, Trisomy 15 |
| 4 | 2 | NB07 | 54840 | 24421 | 51910 | Normal male | Normal male |

<sup>a</sup> UA, uniquely aligned reads. The first 42,500 reads from each samples in run1, and the first 55,000 reads in each sample in run2 were used subjected to aligner to generate UA reads.
<sup>b</sup> Sample 6 is a female, Trisomy 13, Monosomy 14, and large CNV gain on chr 15 (78,914,003-101,997,386) as detected by VeriSeq. The large CNV gain was not significant using 30K reads; it was significant using 60K UA reads.
<sup>c</sup> The mosaic gain on chr6 was not significant using 30K reads; it was significant using 60K UA reads.

**Figure 2:**
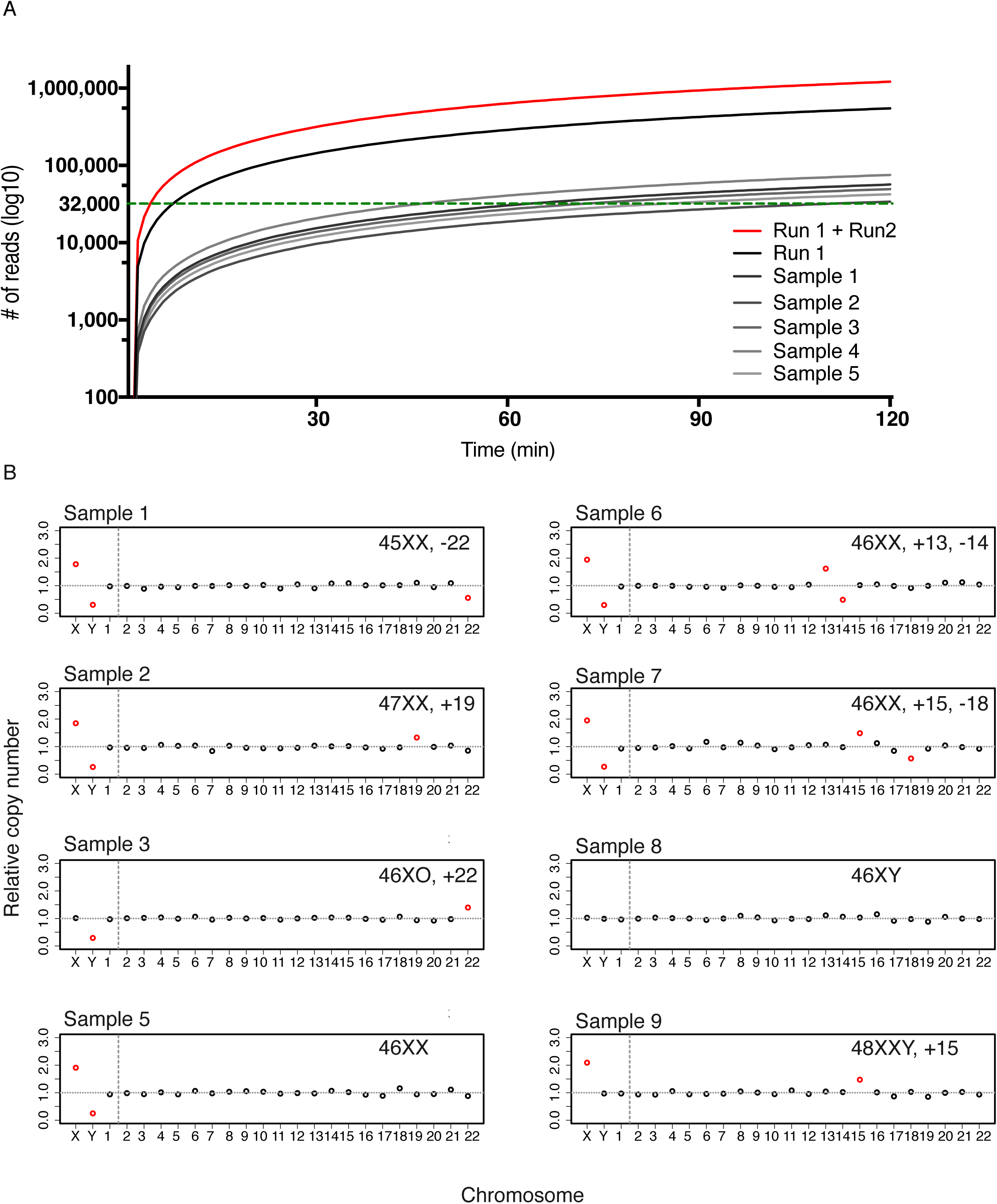
MinION assay results. A). MinION nanopore sequencing representative yield performance (Grayscales: sample 1-5; black: run 1 accumulative; red: run 1 and run 2 accumulative; Green dotted line: threshold for ≥ 30K uniquely aligned reads for downstream analysis). B). MinION-assisted aneuploidy detection results. The abnormality of each chromosome was indicated by color (Red: significantly abnormal; Black: normal).

### Aneuploidy detection

Aneuploidy detection analyses were performed on the basis of Z-score method using the UA reads from each sample generated within 2-hour nanopore sequencing. Based on the known bias introduced from whole genome amplification (WGA), more reads are needed for aneuploidy detection assay when comparing to aneuploidy detection assay on samples that did not undergo WGA. Using the Z-score based assay with incremental read-intervals of 10,000 reads, we found that for whole aneuploidy detection a threshold of 30,000 reads was required. In Run 1, 48-112 minutes sequencing time was needed to generate 30,000 UA reads while in Run 2, 45-98 minutes were needed to generated 30,000 UA reads (Figure 2A, Supple. Figure 1) when 5 samples were included in the same run. If the run time instead of cost is a primary concern and one flow cell is being used dedicatedly for one sample, sufficient reads can be generated for the test in less than 15 minutes (Figure 2A). In our tests, aneuploidy detection results using 30,000-60,000 UA sequencing reads were in agreement with results obtained using VeriSeq NGS tests for whole-chromosome aneuploidy detection (Figure 2B, Table 1). The full trisomy and monosomy cases were detected using 30K or more UA reads and matched with the VeriSeq NGS test results (Figure 2B, Table 1).

### Large CNV detection

The capability of performing large CNV detection using nanopore sequencing was primarily investigated using 60,000 UA reads on sample 6. Sample 6 is a Female with trisomy 13, monosomy 14, and gain on chr15 73M-110M. A large CNV gain on chr 15 (70M-90M) in sample 6 were detected using 60,000 UA nanopore sequencing reads. This shows the potential of performing large CNV detection using nanopore sequencing. However, future experiments are needed to systemically investigate and improve the detection sensitivity and limitation (Supple. Figure 2A).

### Mosaicism detection

Our nanopore-based method was primarily developed to perform rapid aneuploidy screening. However, this method can, in theory, also screen high-level mosaic aneuploidy. Current VeriSeq NGS were developed to screen-out 20-80% mosaic aneuploidy using 600K – 900K reads, representing 1-4 cells carrying full aneuploidy in a 5-cell biopsy. Sample 7 is a female trisomy 15, monosomy 18, and also carries a 20-40% mosaic trisomy on chr6 as determined by VeriSeq NGS. When performing aneuploidy detection using 60,000 UA reads, the trisomy 15 and monosomy 18 were detected, and mosaic trisomy chr6 were detected (Table 1, Supple. Figure 2B). However, the mosaic trisomy on chr6 was not detected when using 30,000 UA reads (Table 1). Thus, it is possible to detect mosaic aneuploidy using nanopore sequencing, but it also requires longer run times in order to generate deeper coverage for high confidence.

### Study limitations

A 1 to 2-hour sequencing run generated sufficient data to perform full aneuploidy detection for up to 5 samples (including a reference control) but is not sufficient for reliable detection of large CNVs or mosaic aneuploidy detection. For that, 2-hour or longer sequencing runs may be required at current sequencing speed, or fewer samples can be run in a single flow cell. In addition, as with other NGS-based approaches, this method would not be able to detect triploid or tetraploid cases when the sex chromosomes are in the same ratio as a normal sample, unless sequencing depth was increased significantly to enable SNP-based screening. Finally, this approach also relies on uniform amplification of the whole genome as an amplification failure can result in noisy result; hence PGS detection failure.

## Discussion

Here we present the first successful application of nanopore sequencing technology (third generation NGS sequencing) to pre-implantation genetic screening (PGS). Our barcoded library preparation of short DNA fragments enabled rapid simultaneous copy number assessment of all chromosomes from four samples on a single relatively low-cost hand-held DNA sequencer, and could be scaled by running multiple nanopore sequencers in parallel. Compared with existing next-generation sequencing technologies, the practical implication of a nanopore-based technique is that PGS testing can be done within an IVF laboratory without the need for expensive equipment or use of extensive space (21, 22). The rapid library preparation and sequencing facilitates same day test-and-transfer of euploid embryos and therefore avoids the need to freeze all embryos and delay transfer waiting for PGS results. In theory, nanopore-based sequencing could also allow simultaneous assessment of aneuploidy (PGS) as well as detection of a predetermined point mutation (PGD) if the WGA-amplified and targeted-amplified DNA were sequenced together, but this would need to be tested and validated.

In contrast to the 2^nd^ generation NGS platforms, such as those from Illumina and Ion Torrent, nanopore-based sequencing delivers sequencing results on a read-by-read basis in real time (21, 22). By reducing the number of samples run on a single nanopore flow-cell or by increasing the number of flow cells used, or sequencing run time, the number of reads on each sample could also be increased further. Increasing the number of reads obtained would allow successful detection of smaller CNVs and mosacism. For example, to detect a 20 mb duplication or deletion, ∼ 60,000 reads would be required. Since the detection of aneuploidy is based on the number of reads aligning to each individual chromosome, the sequencing time necessary for detecting aneuploidy on individual chromosome, particularly of the larger chromosomes such as Chr1 and Chr2, would be even shorter than the time necessary to confirm that the sample is euploid on all chromosomes.

In viewing the results of this study, caution is required. This pilot study was performed on a small number of samples for demonstration purpose, and all samples had been successfully tested by VeriSeq assay. Future prospective studies will be needed to test in larger sample size, and prior to or in parallel with current NGS-or microarray-based PGS testing in order to better estimate the test performance statistics (23). Nonetheless, in the evolution of methods to better assess chromosomal copy number in the preimplantation embryo, from FISH to microarray and NGS (24-26), nanopore-based sequencing has the potential to offer rapid testing results within an IVF lab setting.

## Conclusion

Nanopore-based sequencing is a promising sequencing platform that can enable onsite rapid PGS detection onsite. With a 2.5-hour standard WGA amplification from cell-biopsy, 45 minute library preparation workflow, and < 2-hour sequencing run, normal samples and samples with full aneuploidy were detected correctly. With 2-hour or longer sequencing run, large CNVs and mosaic aneuploidy are feasible.

## Acknowledgement

We sincerely thank Drs. Thomas Tuschl, Jan Vijg, Yousin Suh, members of Center for Women’s Reproductive Care at Columbia University, and members of the Williams lab for their helpful inputs and advices with this project and manuscript. This research was supported by the National Institutes of Health Grant HD068546 and U19CA179564.

